# Genomic Surveillance of *Acinetobacter baumannii* in the Philippines, 2013-2014

**DOI:** 10.1101/2021.03.15.435482

**Authors:** Jeremiah Chilam, Silvia Argimón, Marilyn T. Limas, Melissa L. Masim, June M. Gayeta, Marietta L. Lagrada, Agnettah M. Olorosa, Victoria Cohen, Lara T. Hernandez, Benjamin Jeffrey, Khalil Abudahab, Charmian M. Hufano, Sonia B. Sia, Matthew T.G. Holden, John Stelling, David M. Aanensen, Celia C. Carlos, on behalf of the Philippines Antimicrobial Resistance Surveillance Program

**Affiliations:** Antimicrobial Resistance Surveillance Reference Laboratory, Research Institute for Tropical Medicine, Muntinlupa, Philippines; Centre for Genomic Pathogen Surveillance, Wellcome Genome Campus, Hinxton, UK; University of St Andrews School of Medicine, St Andrews, UK; Brigham and Women’s Hospital, Boston, MA, USA; Big Data Institute, University of Oxford, Oxford, UK

## Abstract

*Acinetobacter baumannii* is an opportunistic nosocomial pathogen that has increasingly become resistant to carbapenems worldwide. In the Philippines, carbapenem resistance and multi-drug resistance (MDR) rates are above 50%. We undertook a genomic study of carbapenem resistant *A. baumannii* in the Philippines to characterize the population diversity and antimicrobial resistance (AMR) mechanisms.

We sequenced the whole genomes of 117 *A. baumannii* isolates recovered by 16 hospitals in the Philippines between 2013 and 2014. We determined the multi-locus sequence type (MLST), presence of acquired AMR determinants and relatedness between isolates from the genome sequences. We also compared the phenotypic and genotypic resistance results.

Carbapenem resistance was mainly explained by the acquisition of class-D beta-lactamase gene *bla*_OXA-23_. The concordance between phenotypic and genotypic resistance to imipenem was 98.15% and 94.97% overall for the seven antibiotics analysed. Twenty-two different sequence types (ST) were identified, including 7 novel STs. The population was dominated by high-risk international clone 2 (i.e., clonal complex 92), in particular by ST195 and ST208 and their single locus variants. With WGS we identified local clusters representing potential undetected nosocomial outbreaks, as well as multi-hospital clusters indicating inter-hospital transmission. Comparison with global genomes suggested that the establishment of carbapenem-resistant IC2 clones in the Philippines is likely the result of clonal expansion and geographical dissemination and at least partly explained by inadequate hospital infection control and prevention.

This study is the first extensive genomic study of carbapenem-resistant *A. baumannii* in the Philippines and underscores the importance of hospital infection control and prevention to contain high-risk clones.

## Introduction

*Acinetobacter baumannii* is one of the most challenging hospital-acquired infections due to its ability to acquire resistance to different groups of antimicrobials and to survive long periods of time on dry surfaces, making eradication in healthcare facilities difficult once it has become endemic. ^1^ A previous surveillance study in the Asia-Pacific region showed that *Acinetobacter* spp. was the most frequently isolated organism from ventilator-associated pneumonia, ^2^ while in recent years the Philippine Antimicrobial Resistance Surveillance Program (ARSP) has been consistently reporting *A. baumannii* as the second and third most commonly isolated organism from cerebrospinal fluid and respiratory specimens, respectively. ^3^

Over the past two decades, *A. baumannii* has become increasingly resistant to carbapenems worldwide, with resistance rates of >40% reported across several Asia-Pacific countries, the highest prevalence of carbapenem resistance amongst important nosocomial gram-negative pathogens. ^4, 5^ This trend is also observed in the Philippines, where the annual resistance rates for several antibiotics, including carbapenems, have been increasing, reaching 56 and 57% in 2017 for meropenem and imipenem, respectively (Figure 1A–C). In addition, the ARSP has reported multi-drug resistant (MDR) rates of 63% and 47% for all isolates and blood isolates, respectively, with combined resistance to aminoglycosides, fluoroquinolones, carbapenems and ampicillin-sulbactam. ^6^ Importantly, bacteremia due to MDR *A. baumannii* has been shown to result in additional hospitalization and costs, compared with bacteremia due to non-MDR *A. baumannii*. ^7^

**Figure 1.**
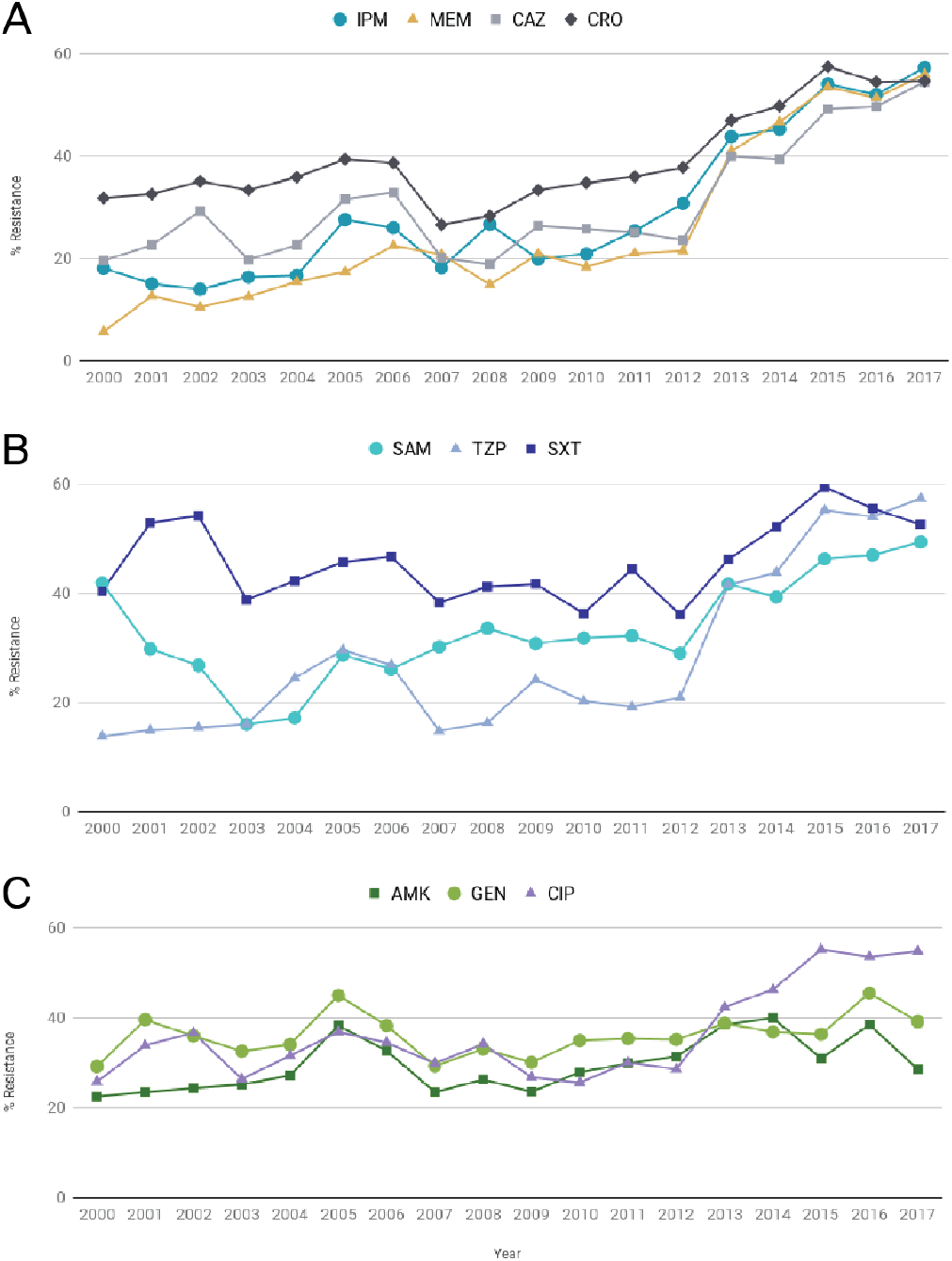
Annual resistance rates of *A. baumannii* between 2000 and 2017. **A)** IPM: imipenem; MEM: meropenem; CAZ: ceftazidime; CRO: ceftriaxone. **B)** SAM: ampicillin-sulbactam; TZP: piperacillin-tazobactam; SXT: sulfamethoxazole-trimetoprim. **C)** AMK: amikacin; GEN: gentamicin; CIP: ciprofloxacin.

Molecular typing methods have shown that clinical isolates of *A. baumannii* with an MDR phenotype belong mostly to two globally disseminated lineages, global clone (GC) 1 and GC2, also known as International Clones (IC)1 and 2. Clonal complex 92 (CC92), corresponding to GC2, was the most prevalent in a previous study across nine Asian countries and including two isolates from the Philippines. ^8^

The ARSP concentrates on phenotypic detection methods for bacterial identification and antimicrobial susceptibility testing. A good understanding on the molecular epidemiology and resistance mechanisms of *A. baumannii* in the country may aid in the control of AMR spread by monitoring the presence of international clones and the emergence of novel lineages. Whole-genome sequencing (WGS) can provide information on antimicrobial resistance and genotyping using a single assay, with added resolution to aid outbreak investigations. ^9^ The current report aims to gather baseline data on the molecular epidemiology of *A. baumannii* with a focus on the predominant circulating lineages and antimicrobial resistance mechanisms.

## Methods

### Bacterial Isolates

A total of 5254 *A. baumannii* isolates were collected and tested for resistance by the ARSP during the period of January 2013 to December 2014. Isolates found to be resistant to carbapenems were subsequently referred to the Antimicrobial Resistance Surveillance Reference Laboratory (ARSRL) for confirmation. Out of the 455 *A. baumannii* isolates referred to the ARSRL in 2013 and 2014 (155 and 290, respectively), a total of 117 isolates representing 16 sentinel sites were selected for whole-genome sequencing (Table 1) according to the following criteria described in detail previously: ^10^ (1) referred to ARSRL in 2013–2014; (2) complete antimicrobial susceptibility data (resistance profile); (3) overall prevalence of the resistance profile in the ARSP data (including both referred and non-referred isolates); (4) geographical representation of different sentinel sites; (5) invasive isolates (i.e., from blood, or cerebrospinal, joint, pleural and pericardial fluids) were selected when both invasive and non-invasive isolates were available for a combination of resistance profile, sentinel site and year of collection. We utilized a proxy definition for “infection origin” whereby patient first isolates collected in the community or on either of the first two days of hospitalization were categorized as community-acquired infection isolates (CA), while isolates collected on hospital day three or later were categorized as hospital-acquired (HA) infection isolates. ^11^

**Table 1.**
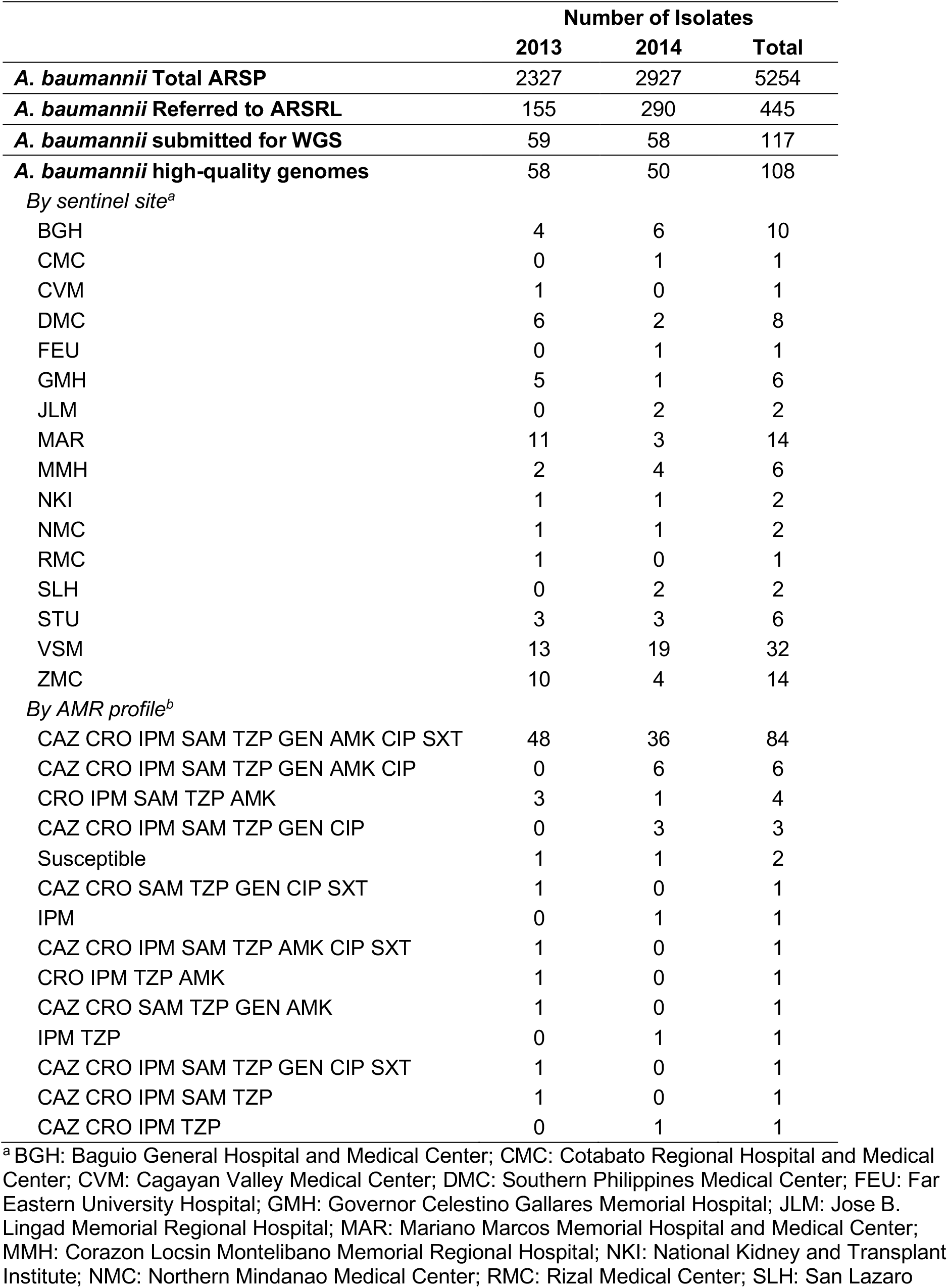
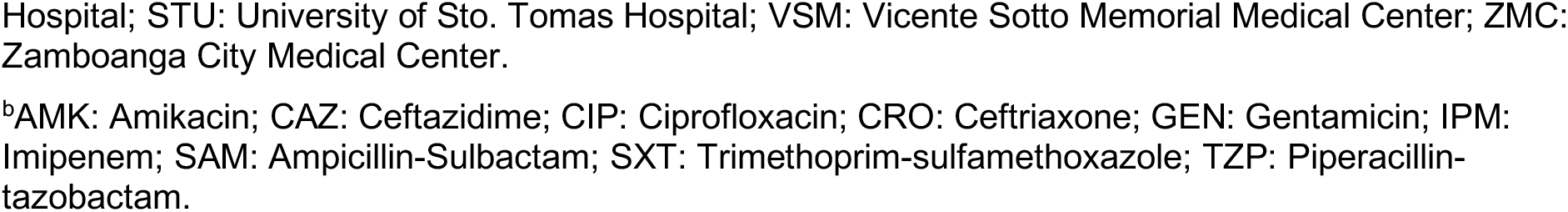
Total number of *A. baumannii* isolates analyzed by the Antimicrobial Resistance Surveillance Program (ARSP) and referred to the Antimicrobial Resistance Surveillance Reference Laboratory (ARSRL) during 2013 and 2014, isolates submitted for whole-genome sequencing, and high-quality *A. baumannii* genomes obtained, discriminated by sentinel site and AMR profile.

|  | Number of Isolates |  |  |
| --- | --- | --- | --- |
|  | 2013 | 2014 | Total |
| ***A. baumannii* Total ARSP** | 2327 | 2927 | 5254 |
| ***A. baumannii* Referred to ARSRL** | 155 | 290 | 445 |
| ***A. baumannii* submitted for WGS** | 59 | 58 | 117 |
| ***A. baumannii* high-quality genomes** | 58 | 50 | 108 |
| *By sentinel site<sup>a</sup>* |  |  |  |
| BGH | 4 | 6 | 10 |
| CMC | 0 | 1 | 1 |
| CVM | 1 | 0 | 1 |
| DMC | 6 | 2 | 8 |
| FEU | 0 | 1 | 1 |
| GMH | 5 | 1 | 6 |
| JLM | 0 | 2 | 2 |
| MAR | 11 | 3 | 14 |
| MMH | 2 | 4 | 6 |
| NKI | 1 | 1 | 2 |
| NMC | 1 | 1 | 2 |
| RMC | 1 | 0 | 1 |
| SLH | 0 | 2 | 2 |
| STU | 3 | 3 | 6 |
| VSM | 13 | 19 | 32 |
| ZMC | 10 | 4 | 14 |
| *By AMR profile<sup>b</sup>* |  |  |  |
| CAZ CRO IPM SAM TZP GEN AMK CIP SXT | 48 | 36 | 84 |
| CAZ CRO IPM SAM TZP GEN AMK CIP | 0 | 6 | 6 |
| CRO IPM SAM TZP AMK | 3 | 1 | 4 |
| CAZ CRO IPM SAM TZP GEN CIP | 0 | 3 | 3 |
| Susceptible | 1 | 1 | 2 |
| CAZ CRO SAM TZP GEN CIP SXT | 1 | 0 | 1 |
| IPM | 0 | 1 | 1 |
| CAZ CRO IPM SAM TZP AMK CIP SXT | 1 | 0 | 1 |
| CRO IPM TZP AMK | 1 | 0 | 1 |
| CAZ CRO SAM TZP GEN AMK | 1 | 0 | 1 |
| IPM TZP | 0 | 1 | 1 |
| CAZ CRO IPM SAM TZP GEN CIP SXT | 1 | 0 | 1 |
| CAZ CRO IPM SAM TZP | 1 | 0 | 1 |
| CAZ CRO IPM TZP | 0 | 1 | 1 |

<sup>a</sup> BGH: Baguio General Hospital and Medical Center; CMC: Cotabato Regional Hospital and Medical Center; CVM: Cagayan Valley Medical Center; DMC: Southern Philippines Medical Center; FEU: Far Eastern University Hospital; GMH: Governor Celestino Gallares Memorial Hospital; JLM: Jose B. Lingad Memorial Regional Hospital; MAR: Mariano Marcos Memorial Hospital and Medical Center; MMH: Corazon Locsin Montelibano Memorial Regional Hospital; NKI: National Kidney and Transplant Institute; NMC: Northern Mindanao Medical Center; RMC: Rizal Medical Center; SLH: San Lazaro
Hospital; STU: University of Sto. Tomas Hospital; VSM: Vicente Sotto Memorial Medical Center; ZMC: Zamboanga City Medical Center.
<sup>b</sup>AMK: Amikacin; CAZ: Ceftazidime; CIP: Ciprofloxacin; CRO: Ceftriaxone; GEN: Gentamicin; IPM: Imipenem; SAM: Ampicillin-Sulbactam; SXT: Trimethoprim-sulfamethoxazole; TZP: Piperacillin-tazobactam.

### Antimicrobial Susceptibility Testing (AST)

All *A. baumannii* isolates from this study were tested for antimicrobial susceptibility to nine antibiotics representing six different classes, namely ceftazidime (CAZ), ceftriaxone (CRO), imipenem (IPM), ampicillin-sulbactam (SAM), piperacillin-tazobactam (TZP), gentamicin (GEN), amikacin (AMK), ciprofloxacin (CIP), and trimethoprim-sulfamethoxazole (SXT) (Table 1). Antimicrobial susceptibility of the isolates was determined at the ARSRL using one or a combination of the following methods, Kirby-Bauer disk diffusion method, gradient methods such as E-Test and/or Vitek 2 Compact automated system (BioMérieux, Marcy-l’Étoile, France). The zone of inhibition and minimum inhibitory concentration obtained were interpreted according to the 26th edition of the Clinical and Laboratory Standard Institute (CLSI) guidelines ^12^ to determine the resistance profile of the isolates as the list of antimicrobials to which the organism is non-susceptible. Multi-drug resistant (MDR) and extensively drug resistant (XDR) phenotypes were classified as per standard definitions. ^13^

### DNA Extraction and Whole-Genome Sequencing

DNA was extracted from a single colony of each of 117 *A. baumannii* isolates with the QIAamp 96 DNA QIAcube HT kit and a QIAcube HT (Qiagen; Hilden, Germany). DNA extracts were multiplexed and sequenced on the Illumina HiSeq platform (Illumina, CA, USA) with 100-bp paired-end reads. Raw sequence data were deposited in the European Nucleotide Archive (ENA) under the study accession PRJEB17615. Run accessions are provided on the Microreact projects.

### Bioinformatics analysis

Genome quality was assessed based on metrics produced for assemblies, annotation files, and the alignment of the reads to the reference genome *A. baumanni* strain ATCC 17978 (accession CP000521), as previously described. ^10^ Annotated assemblies were produced from short-read Illumina data as previously described. ^14^ We derived *in silico* the multi-locus sequence type (MLST) of the isolates from the whole genome sequences. The sequence types (ST) were determined from assemblies with Pathogenwatch (https://pathogen.watch/) or from sequence reads with ARIBA ^15^ and the *A. baumannii* database hosted at PubMLST. ^16^ The isolates were assigned to international clones (IC) based on their ST as previously indicated. ^17–20^

Evolutionary relationships between isolates were inferred from single-nucleotide polymorphisms (SNPs) by mapping the paired-end reads to the reference genomes of *A. baumannii* strains A1 (accession CP010781) or AC29 (ST195, CC92, accession CP007535), as described in detail previously. ^10^ Mobile genetic elements (MGEs) were masked in the alignment of pseudogenomes with a script available at https://github.com/sanger-pathogens/remove_blocks_from_aln. Alignments of SNP positions were inferred with snp-sites v2.4.1. ^21^ For the phylogenies of CC92 genomes, recombination regions detected with Gubbins ^22^in the alignment of pseudogenomes were also removed. Maximum likelihood phylogenetic trees were generated with RAxML v8.28 ^23^ based on the generalised time reversible (GTR) model with GAMMA method of correction for among-site rate variation and 100 bootstrap replications. Pairwise SNP differences between primary isolates belonging to the same or to different hospitals were calculated from alignments of SNP positions with a script available at https://github.com/simonrharris/pairwise_difference_count.

To contextualize the Philippine genomes, global *A. baumannii* genomes with geolocation and isolation date mainly between 2007 and 2017 for which raw Illumina paired-end sequence data were available at the European Nucleotide Archive were downloaded, assembled and quality controlled as above. Evolutionary relationships between global genomes and those from this study were inferred from an alignment of SNP positions obtained after mapping the reads to the complete genome of strain A1 and masking regions with mobile genetic elements as described above. The tree of 977 genomes was obtained using an approximately-maximum likelihood phylogenetic method with FastTree ^24^. The tree of 573 global CC92 genomes was inferred with RAxML from an alignment of SNP sites obtained after mapping the genomes to the complete genome of strain AC29 and removing mobile genetic elements and recombination regions, as described above.

Known AMR determinants were identified from raw sequence reads using ARIBA ^15^ and two different AMR databases, a curated database of acquired resistance genes ^25^, and the Comprehensive Antibiotic Resistance Database (CARD ^26^). Point mutations were identified on gyrase and topoisomerase genes with CARD and ARIBA, and corroborated with a literature search. The presence of the insertion sequences IS*Aba1* (accession AY758396) and IS*Aba125* (accession AY751533) upstream of the *ampC* gene was examined with ISMapper v 2.0.1 ^27^ using the reference genome of *A. baumannii* A1 (accession CP010781) and default parameters. Genomic predictions of resistance were derived from the presence of known antimicrobial resistance genes and mutations identified in the genome sequences. The genomic predictions of AMR (test) were compared to the phenotypic results (reference) and the concordance between the two methods was computed for each of 7 antibiotics (756 total comparisons). Isolates with either a resistant or an intermediate phenotype were considered non-susceptible for comparison purposes. An isolate with the same outcome for both the test and reference (i.e., both susceptible or both non-susceptible) was counted as a concordant isolate. The concordance was the number of concordant isolates over the total number of isolates assessed (expressed as percent).

All project data, including inferred phylogenies, AMR predictions and metadata were made available through the web application Microreact (http://microreact.org).

## Results

### Demographic and Clinical Characteristics of the *A. baumannii* Isolates

Out of the 117 *A. baumannii* genomes sequenced (Table 1), 7 genomes were excluded based on quality, while 2 genomes were identified *in silico* as *Acinetobacter pitti*. The demographic and clinical characteristics of the remaining 108 *A. baumannii* isolates with high-quality genomes are summarized on Table 2. The age of the patients ranged from less than 1 year to 92 years old, with 31% of the isolates (*n*=34) from patients 65 years old and above. Sixty-two per cent of the isolates (*n*=67) were from males. The majority of the isolates (99.1%, *n*=107) were from in-patients, and classified as from a hospital-acquired infection (76.85%, *n*=83). Respiratory samples (tracheal aspirates and sputum) accounted for 55.56% of the specimens (*n*=60).

**Table 2.** Demographic and clinical characteristics of 108 sequenced and confirmed *A. baumannii* isolates collected from 16 ARSP sites.

| Characteristic | No. Isolates |
| --- | --- |
| **Sex** |  |
| Male | 67 |
| Female | 41 |
| **Age (in years)** |  |
| <1 | 6 |
| 1-4 | 11 |
| 5-14 | 3 |
| 15-24 | 6 |
| 25-34 | 7 |
| 35-44 | 9 |
| 45-54 | 12 |
| 55-64 | 20 |
| 65-80 | 26 |
| >=81 | 8 |
| **Patient Type** |  |
| In-patient | 107 |
| Out-patient | 1 |
| **Specimen Origin** |  |
| Community-acquired | 25 |
| Hospital-acquired | 83 |
| **Submitted As\*** |  |
| Carbapenem-resistant | 104 |
| Non carbapenem-resistant | 4 |
| **Specimen Type** |  |
| Aspirate | 1 |
| Blood** | 21 |
| Bone | 1 |
| Catheter | 1 |
| Catheter, central | 1 |
| Cerebrospinal fluid** | 13 |
| Sputum | 10 |
| Tracheal aspirate | 50 |
| Ulcer | 1 |
| Urine | 4 |
| Wound | 5 |

\* Specimen Origin is computed based on admission date of the patient
\*\* Specimen types considered as Invasive isolates.

### Concordance between phenotypic and genotypic AMR results

The genotypic predictions of AMR were highly concordant with the phenotypic results (overall concordance was 94.97%, Table 3). The concordance for imipenem was 98.15% and, of the 104 resistant isolates, 97 isolates from 14 hospitals (93.26%) carried the class D beta-lactamase gene *bla*_OXA-23_, alone or in combination with *bla*_OXA-235_ (*n*=1). The remaining isolates carried *bla*_NDM-6_ (*n*=3), *bla*_NDM-1_ (*n*=2) or *bla*_OXA-72_ (*n*=2). One isolate had no known acquired carbapenemase. Of the 104 isolates resistant to imipenem 89 (85.58%) were classified as XDR and 13 (12.50%) as MDR, with the notable presence of the *armA* gene encoding a 16S rRNA methyltransferase conferring broad-spectrum resistance to aminoglycosides in 54 isolates, and the co-occurrence of mutations in *gyrA* and *parC* conferring resistance to fluoroquinolones in 95 isolates (Table 3). Acquired colistin resistance genes (*mcr*) were not detected.

**Table 3.**
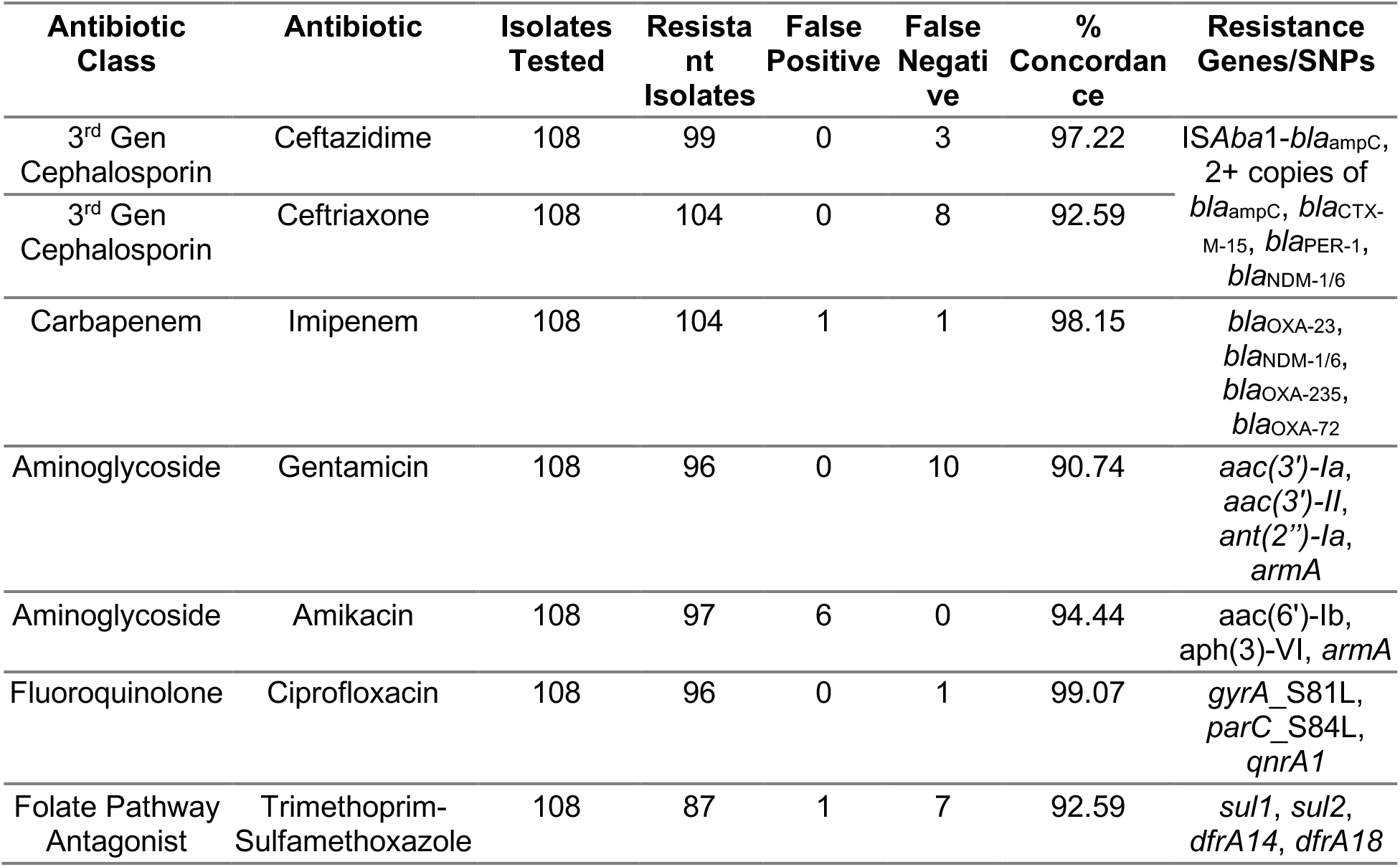
Comparison between antimicrobial susceptibility testing results and genotypic resistance for 108 *A. baumannii* isolates.

| Antibiotic Class | Antibiotic | Isolates Tested | Resistant Isolates | False Positive | False Negative | % Concordance | Resistance Genes/SNPs |
| --- | --- | --- | --- | --- | --- | --- | --- |
| 3<sup>rd</sup> Gen Cephalosporin | Ceftazidime | 108 | 99 | 0 | 3 | 97.22 | ISAba1-*bla*<sub>ampC</sub>, 2+ copies of *bla*<sub>ampC</sub>, *bla*<sub>CTX-M-15</sub>, *bla*<sub>PER-1</sub>, *bla*<sub>NDM-1/6</sub> |
| 3<sup>rd</sup> Gen Cephalosporin | Ceftriaxone | 108 | 104 | 0 | 8 | 92.59 |  |
| Carbapenem | Imipenem | 108 | 104 | 1 | 1 | 98.15 | *bla*<sub>OXA-23</sub>, *bla*<sub>NDM-1/6</sub>, *bla*<sub>OXA-235</sub>, *bla*<sub>OXA-72</sub> |
| Aminoglycoside | Gentamicin | 108 | 96 | 0 | 10 | 90.74 | *aac*(3')-Ia, *aac*(3')-II, *ant*(2'')-Ia, *armA* |
| Aminoglycoside | Amikacin | 108 | 97 | 6 | 0 | 94.44 | *aac*(6')-Ib, *aph*(3)-VI, *armA* |
| Fluoroquinolone | Ciprofloxacin | 108 | 96 | 0 | 1 | 99.07 | *gyrA*_S81L, *parC*_S84L, *qnrA1* |
| Folate Pathway Antagonist | Trimethoprim-Sulfamethoxazole | 108 | 87 | 1 | 7 | 92.59 | *sul1*, *sul2*, *dfrA14*, *dfrA18* |

Isolates non-susceptible to third generation cephalosporins ceftazidime (*n*=99) and/or ceftriaxone (*n*=104) carried either the insertion sequence IS*Aba1* upstream of the chromosomal *bla*_ampC_ gene (*n*=67), two or three copies of the *bla*_ampC_ gene (*n*=22), the extended-spectrum beta-lactamase (ESBL) genes *bla*_PER-1_ (*n*=4) and *bla*_CTX-M-15_ (*n*=1), or the carbapenemase gene *bla*_NDM_ (*n*=5). Most of the false negative calls for ceftazidime (*n*=3) and ceftriaxone (*n*=8, Table 3) for which no resistance mechanism was detected, coincided with intermediate susceptibility (*n*=2 and *n*=5, respectively).

### Genotypic findings

#### In silico *genotyping*

Multi-locus sequence type was predicted *in silico* from the whole-genome sequence data of the 108 *A. baumannii* isolates. A total of 22 different STs were identified from this data set as per the Oxford scheme, seven of which were novel and now identified as ST2197, 2199, 2220, 2317, 2318, 2319 and 2320. The population was dominated by clonal complex (CC)92 (*n*=61), represented mainly by ST195 (*n*=29) and ST208 (*n*=23). CC92 was found in 13 out of the 16 sentinel sites, with ST195 and ST208 spread geographically across 8 and 7 sentinel sites, respectively. In contrast, ST369 (*n*=5) was found in only one site. The *armA* gene was found only in isolates belonging to CC92 (*n*=54) from 11 hospitals. Seven of the eight hospitals represented by six or more sequenced isolates showed clonal diversity, with at least two different circulating STs (Table 4). In the remaining hospital (Baguio General Hospital, BGH) all 10 isolates belonged to the same ST (208).

**Table 4.**
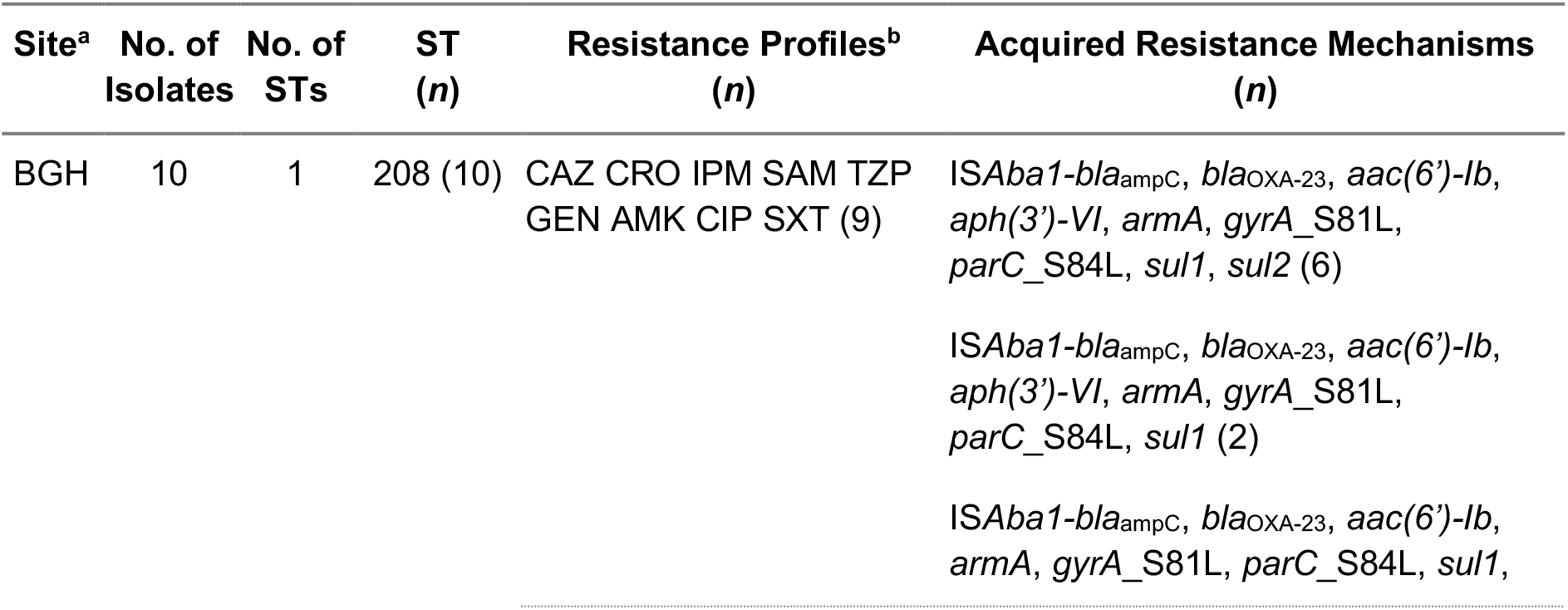
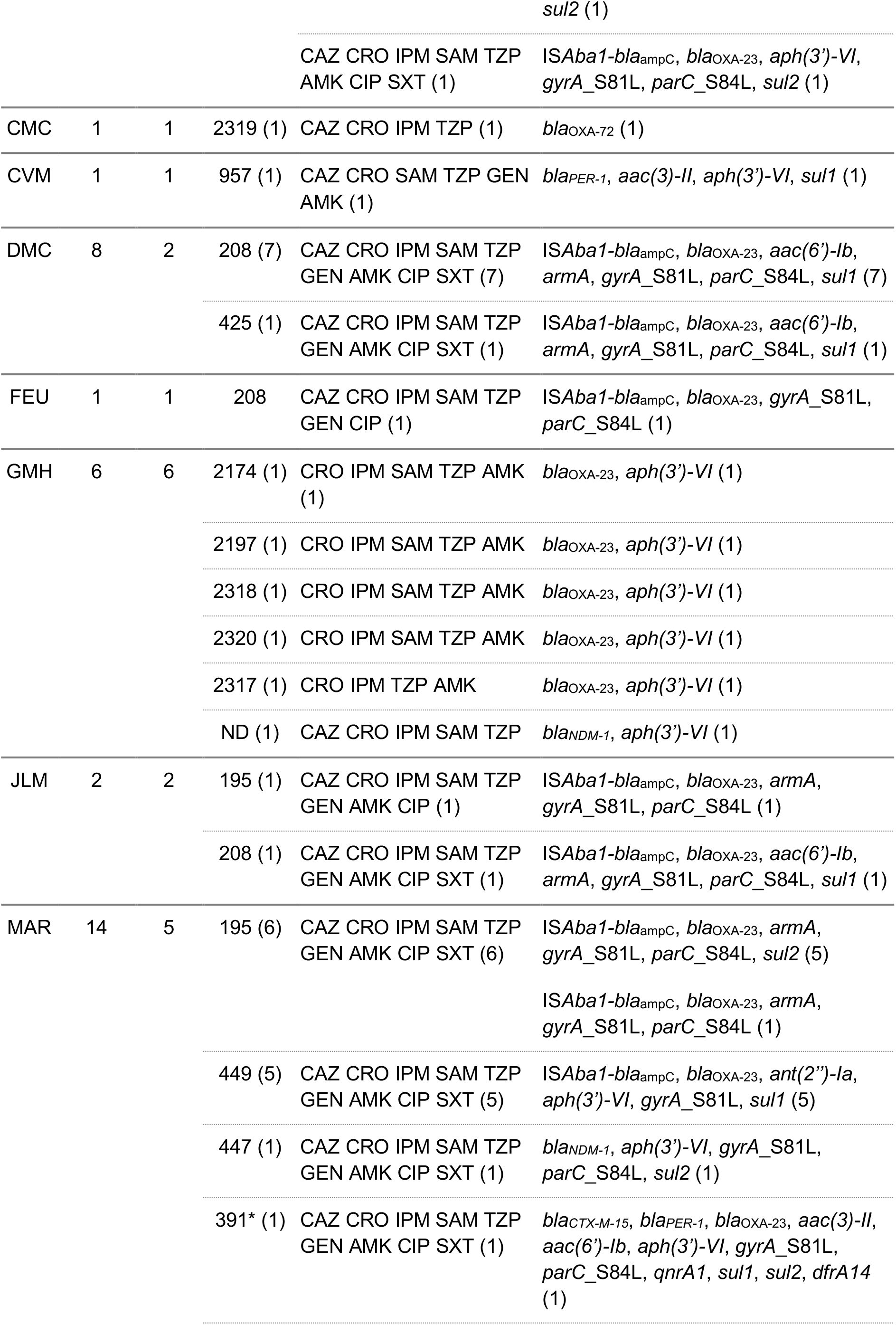
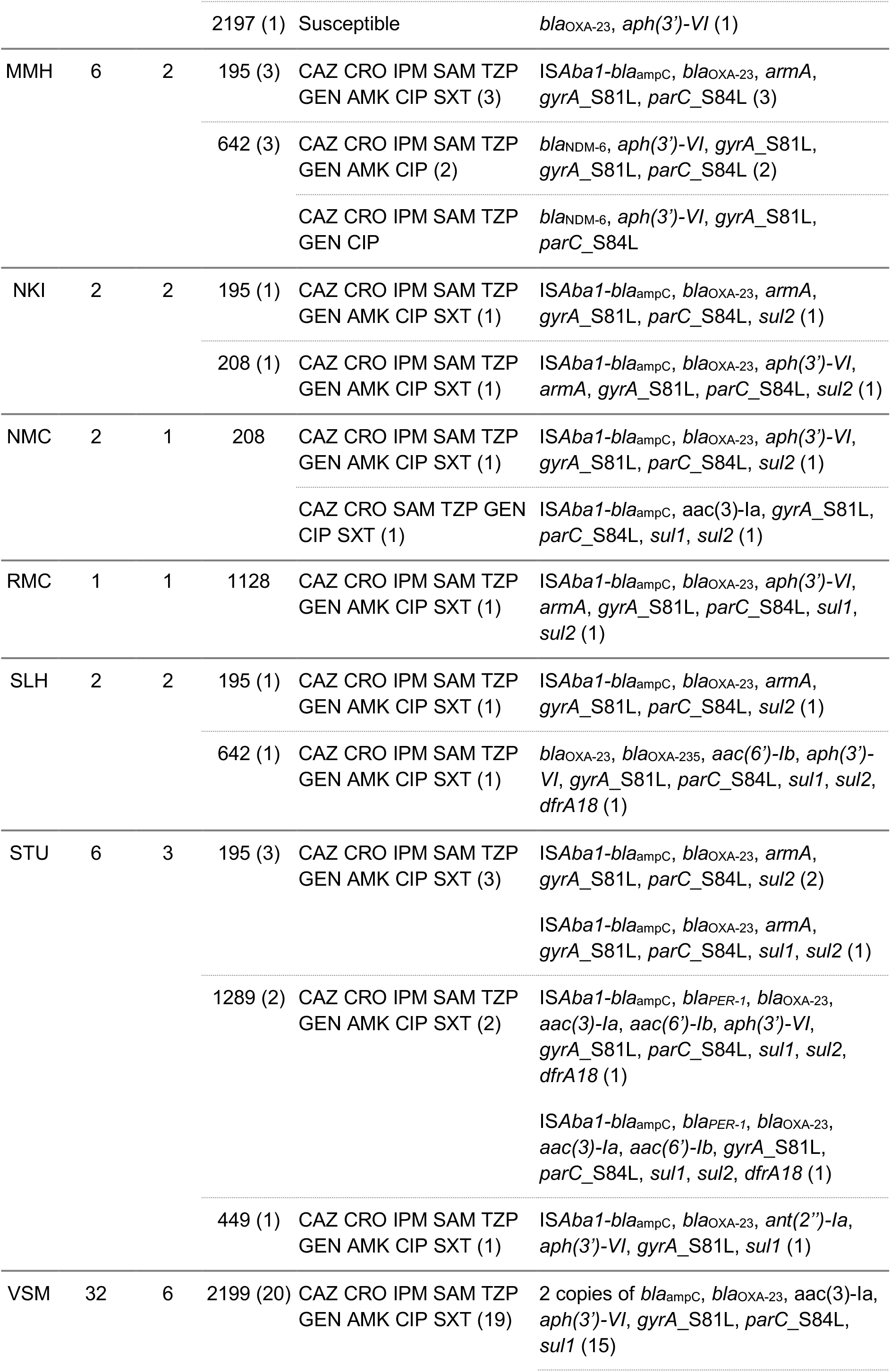
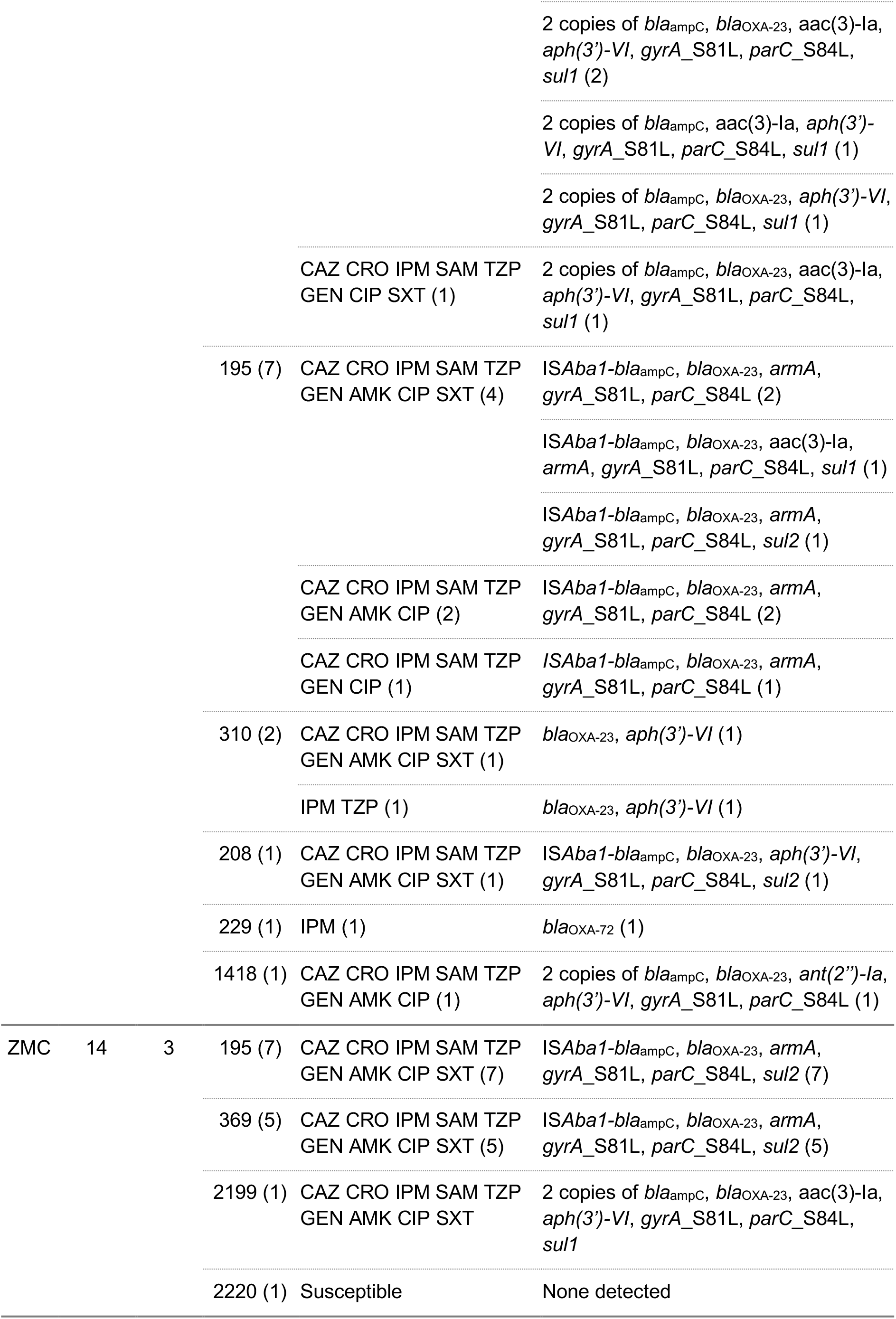
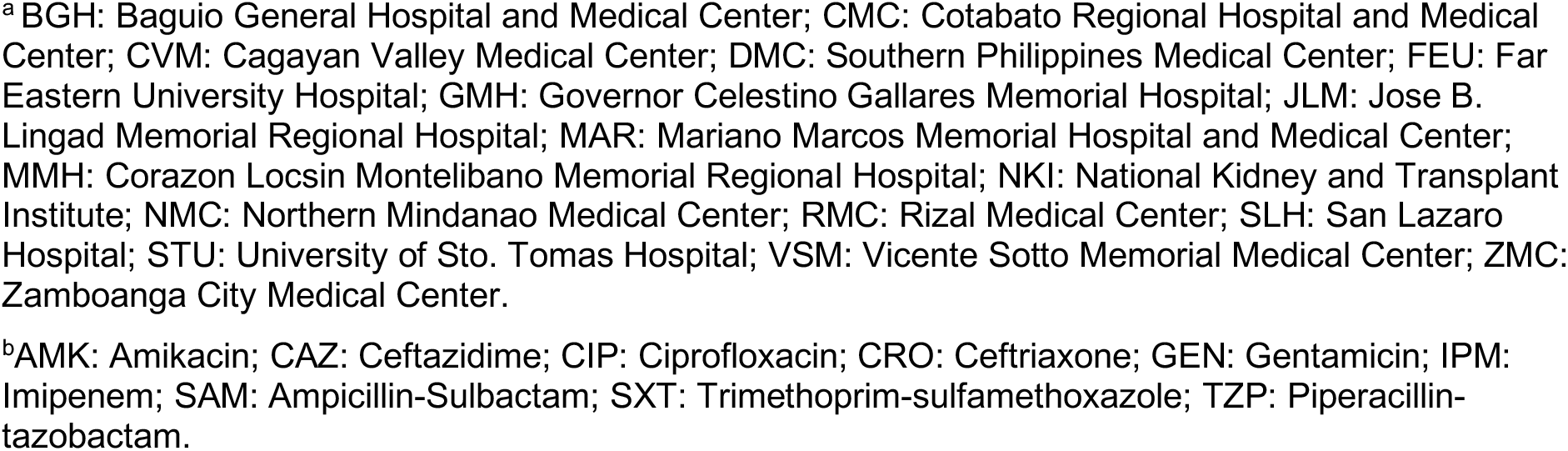
The summary of distribution, sequence types (ST), resistance profiles and Antimicrobial Resistance genes and mutations of the 108 isolates collected from 17 Antimicrobial Resistance Surveillance Program sentinel sites.

#### Population structure of A. baumannii in the Philippines

The phylogenetic tree of 108 *A. baumannii* genomes showed that the population was composed of well-defined clades that matched the distribution of the STs. The two main clonal groups were international clones IC1 and IC2 (i.e., CC91, Figure 2A), with a minor representation of IC8 and IC7. Isolates belonging to international clones were mostly XDR and are known to be responsible for disseminating AMR globally. The carbapenemase gene *bla*_OXA-23_ was found consistently in IC1 and IC2 genomes, and more sporadically in IC8 and non-clonal genomes. In contrast, the carbapenemase gene *bla*_NDM-6_ was found exclusively in three IC8 genomes from Corazon Locsin Montelibano Memorial Regional Hospital (MMH), while *bla*_NDM-1_ and *bla*_OXA-72_ were only found sporadically. Notably, isolates carrying IS*Aba1* inserted in the promoter of *bla*_ampC_ belonged to ST449 (IC1) or to CC92 (IC2), while isolates carrying two or three copies of the *bla*_ampC_ gene all belonged to a novel ST (now ST2199) found in the Vicente Sotto Memorial Medical Center (VSM) in the Visayas region and the Zamboanga Medical Center (ZMC) in the Mindanao region (Figure 2A).

**Figure 2.**
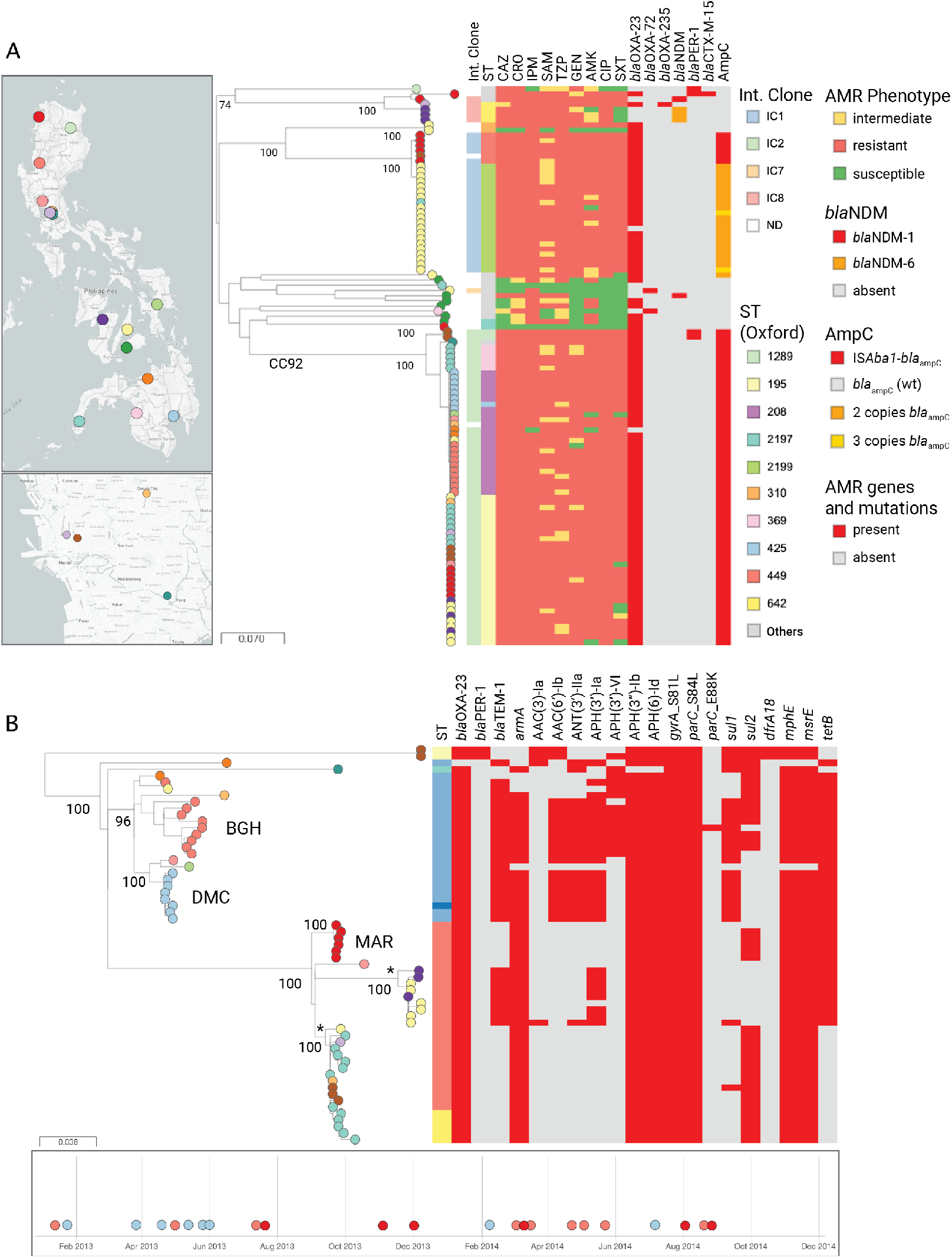
Genomic surveillance of *A. baumannii* from the Philippines 2013–2014. **A)** Phylogenetic tree of 108 isolates inferred from an alignment of 168,916 SNP sites obtained after mapping the genomes to the complete genome of strain A1 and masking MGEs from the alignment. The tree leaves are coloured by sentinel site and indicated on the map (left panels, top: Philippines, bottom: detail of the National Capital Region). The tree is annotated with the isolates assignment to international clones and sequence type, the susceptibility testing results and the presence of acquired carbapenemase genes (ST). The full data are available at https://microreact.org/project/ARSP_ABA_2013-2014. **B)** Phylogenetic tree of 61 CC92 genomes, inferred from an alignment of 618 SNP sites after mapping the genomes to reference AC29 and removing MGEs and recombination regions. The tree leaves are coloured by sentinel site, as indicated on the map from Fig. 2A. The tree blocks represent the distribution of sequence types (STs) and of acquired resistance genes and mutations. Three hospital clusters are annotated on the tree with the hospital code (BGH, DMC, MAR) and their isolation dates are indicated on the timeline. Two multi-hospital clusters are annotated with an asterisk. The full data are available at https://microreact.org/project/ARSP_ABA_CC92_2013-2014. The scale bars represent the number of single nucleotide polymorphisms per variable site.

The phylogenetic tree of 61 genomes from the prevalent XDR CC92 clone showed that most isolates grouped into two clades represented by ST208 and single locus variant (SLV) ST425 (bootstrap support 96%) and by ST195 and SLV ST369 (bootstrap support 100%, Figure 2B). Both ST208-ST425 and ST195-ST369 were found in hospitals from all three island groups (Luzon in the north, Visayas in the center, and Mindanao in the south), but their geographical distributions showed little overlap. The phylogeographic signal suggested both local outbreaks and inter-hospital dissemination (Figure 2B). We investigated this further by counting the number of pairwise, non-recombinant SNP differences between primary isolates from the same or different hospitals. First, we identified three intra-hospital clusters (100% bootstrap support) of closely related isolates from Baguio General Hospital (BGH, ST208, 2-35 pairwise SNPs, *n*=9), Southern Philippines Medical Center (DMC, ST208-425, 1-6 pairwise SNPs, *n*=8), and Mariano Marcos Memorial Hospital and Medical Center (MAR, ST195, 0-3 pairwise SNPs, *n*=6). The isolates within each of the three clusters carried identical or almost identical repertoires of resistance determinants, further supporting their clonal relationship. The isolation dates spanning over 12 months, suggested that these clonal lineages are possibly endemic to the hospitals, although regular introduction by colonized patients cannot be ruled out. Second, we identified two clusters of closely related isolates from two or more hospitals. One cluster contained nine ST195 genomes from two hospitals in the Visayas region (MMH and VSM) with a median of only 5 pairwise SNP differences (range 1-17) between isolates from different hospitals. The second one contained 18 ST195-ST369 genomes from six hospitals across three different regions, with a median of 25 pairwise SNP differences (range 1-53). The clonal relationship between isolates from different hospitals within these two clusters is also supported by a similar complement of resistance determinants.

#### A. baumanni *from the Philippines in global context*

To place the retrospective collection of *A. baumannii* isolates from the Philippines in the context of the global population of this pathogen, we compared our genomes to 931 public genomes available from sequence data archives with linked geographic and temporal information. The isolates were collected between 1982 and 2016, with 94.7% of the isolates collected on 2007 onwards. The public genomes belonged to 16 countries and were assigned to 154 STs in the Oxford scheme. The population represented by the global genomes was substantially skewed towards genomes from the USA (40.5% of the genomes), and belonging to CC92 (58.6%). The Philippine genomes were found in multiple branches of the tree as expected by the diversity of STs, but mostly forming discreet clusters within each branch without genomes from other countries interspersed (Figure 3A). This suggests that the establishment of each clone in the Philippines is the result of one or few founding events.

**Figure 3.**
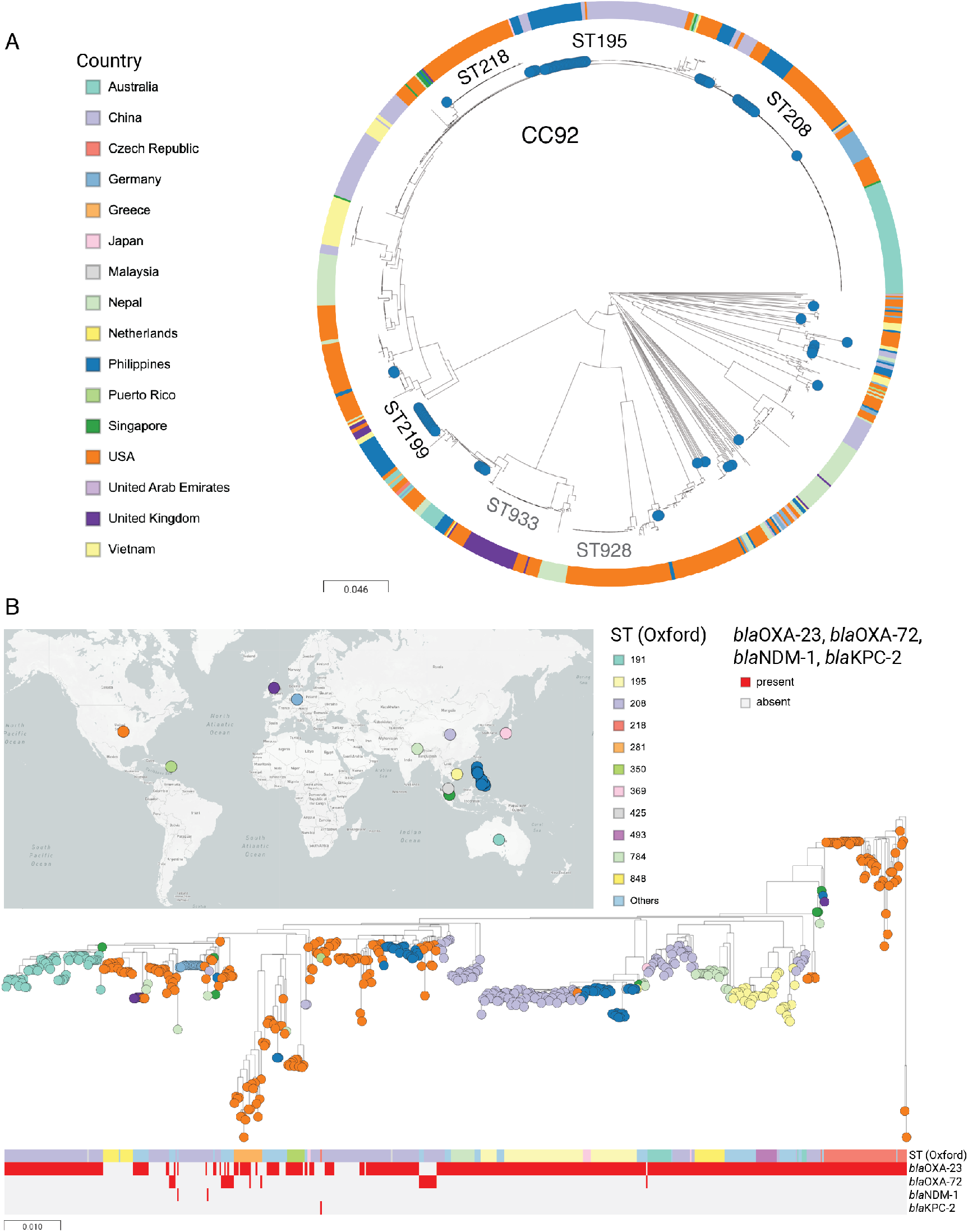
*A. baumannii* from the Philippines in global context. **A)** Phylogenetic tree of 977 isolates from the Philippines (blue nodes) and from 15 other countries inferred from 305,031 SNP positions. The major STs and CCs are labelled in black if represented by genomes of this study, or in grey if they are not. The data are available at https://microreact.org/project/ARSP_ABA_Global. **B)** Phylogenetic tree of 573 CC92 isolates inferred from an alignment of 5,890 SNP positions. The tree leaves are coloured by country as indicated on the map. The tree is annotated with the distribution of acquired carbapenemase genes (red: present, grey: absent). The data are available at https://microreact.org/project/ARSP_CC92_Global. The scale bars represent the number of single nucleotide polymorphisms (SNPs) per variable site.

To investigate in more detail the relationship to global genomes within CC92, a tree of 573 genomes was inferred from the alignment of non-recombinant SNPs. The ST195-ST369 genomes from the Philippines were related to genomes from Singapore, Vietnam, Malaysia, China and USA, while the ST208-ST425 genomes were related to genomes from China, USA and Puerto Rico. However, the strong phylogeographic signal displayed by both the ST195-ST369 and the ST208-ST425 subtrees suggested a single founder event in the Philippines for each clone, followed by their expansion.

## Discussion

In the present study we report on the combined genomic and laboratory-based surveillance of *A. baumannii* in the Philippines during 2013-2014. The prevalence of carbapenem-resistant *A. baumannii* during this period was above 40%, and we therefore focused on the characterization of these organisms. In *A. baumannii*, only low-level carbapenem resistance is mediated by the chromosomal OXA-51-like carbapenemase. The class D OXA-23 carbapenemase was the most prevalent acquired carbapenem resistance mechanism identified in this study, in line with global trends. ^28^ We also detected representatives from the OXA-235-like (*bla*_OXA-235_) and the OXA-40-like groups (*bla*_OXA-72_) albeit in low frequency. No OXA-58-like carbapenemases were detected, as previously reported from other Asia Pacific nations. ^29^ Importantly, we also detected the presence of class B metallo-beta-lactamases NDM-1 and NDM-6 which, unlike OXA-23, confer resistance to extended-spectrum cephalosporins as well as carbapenems. *A. baumannii* harbouring NDM-1 has been sporadically reported previously from other countries ^30–32^, but NDM-6-carrying *A. baumannii* has only recently been reported from Spain. ^33^ Resistance to extended-spectrum cephalosporins was mainly explained by the insertion of IS*Aba1* in the promoter of the intrinsic gene *bla*_ampC_, which has been shown to lead to increased expression of the encoded cephalosporinase. ^34^ Identification of this mechanism represents an additional *in silico* query of the genomes, which is burdensome in the context of a public health reference laboratory, but omitting it would lead to high very major error rates for genomic predictions of resistance to extended-spectrum cephalosporins.

Both IC1 and IC2 were found in the Philippines, both of which are prevalent worldwide and responsible for the spread of MDR and XDR phenotypes. ^28, 35^However, IC2 was the predominant clonal type of *A. baumannii* in our study population, with ST195 and ST208 and respective SLVs found throughout the country. The global phylogenetic tree showed that these two lineages diverged before their establishment in the Philippines. The genetic relatedness of isolates from different hospitals and their similar complement of resistance determinants supports the notion that their subsequent success was the result of clonal expansion and in-country geographic dissemination, rather than multiple introductions, and highlights the need for concerted infection prevention and control measures to contain the spread of high-risk clones. However, the limited number and disparate sampling of genomes from other countries in the region limits our ability to capture the dynamics of these clones.

We also identified three ST195 and ST208 intra-hospital clusters spanning over twelve months each. Resistance to antimicrobial drugs and to desiccation contribute to the survival of *A. baumannii* in the hospital environment, ^1^ and cross-contamination of hospital surfaces with MDR strains has been documented, in particular the areas surrounding colonized or infected patients. ^36, 37^ The ARSP surveillance does not currently include environmental samples and thus it was not possible to connect the persistence of the intra-hospital clusters to environmental contamination. Outbreaks of *A. baumannii* with *bla*_OXA-23_, including of ST195 and ST208, have been reported from several countries, ^38–40^and our study identified potential hospital outbreaks retrospectively. The resolution afforded by WGS was in stark contrast with the uniform resistance profiles of the isolates in our study, thus making cluster detection based on WGS rather than resistance profiles of particular utility for carbapenem-resistant *A. baumannii*.

Assignment of isolates to an outbreak based on their genetic distance is key for effective patient containment and infection control during an ongoing investigation. Out of the three intra-hospital IC2 clusters detected, the ST208 cluster from BGH displayed more genetic diversity than the other two based on the number of pairwise SNP differences, opening the possibility that more than one closely related strains were circulating in the hospital. However, the absence of data on patient movement precluded epidemiological investigation to aid the delineation of outbreaks. In addition, while the pairwise SNP differences are similar to those reported in other studies, ^39, 41–43^SNP thresholds are difficult to assess by comparison, due to methodological differences, such as the use of core- vs whole-genome SNPs, the choice of reference genome for reference-based mapping of short reads, and the inclusion or exclusion of SNPs associated with recombination regions.

In conclusion, our retrospective genomic epidemiology study of carbapenem-resistant *A. baumannii* in the Philippines revealed IC2 with OXA-23 is the main culprit behind the increasing carbapenem resistance rates in the Philippines, and that breaches in infection control and prevention likely contributed to its dissemination. WGS proved a useful tool to improve surveillance of *A. baumannii*.

